# Positive selection of neuregulin 1 (NRG-1) among three long-lived vertebrates

**DOI:** 10.1101/2023.03.18.533285

**Authors:** Neil Copes, Clare-Anne Edwards Canfield

**Affiliations:** Moirai Conservation and Research, Dade City, FL, USA

## Abstract

Maximum lifespan is a species-specific trait that can vary over a broad range, even between closely related species. We selected the few long-lived vertebrates for which reference genome data is available, specifically the blue whale *Balaenoptera musculus* and the Pinta Island tortoise *Chelonoidis abingdonii*. For these species, we used established methods of CpG dinucleotide analysis and allometric estimation to compare predicted maximum longevity with the maximum longevities reported in the AnAge database and in the literature. Additionally, we compared protein sequences between these species and closely related short-lived vertebrates. Orthologous protein sequences with higher pairwise alignment scores between either long-lived species, or between short-lived species, than between long-lived and short-lived relatives were identified. High-scoring orthologs were investigated for evidence of positive selection and convergent evolution, and for evidence of deleterious effect on phenotype. Analysis revealed no evidence for either convergent evolution or predicted deleterious protein sequences changes within these orthologs, however evidence of positive selection was identified in two genes: NRG-1 and GALNT17. Further comparison to protein sequences from the long-lived bowhead whale (*Balaena mysticetus*) additionally supported NRG-1 as exhibiting evidence of positive selection among the selected long-lived species.

## Introduction

Lifespans vary much less within species than they do between species. For example, as of 2013 the average worldwide human life expectancy beyond the age of 10 was 77.4 years, with a standard deviation of only 14.7 years (Edwards 2013). By contrast, the range of maximum lifespans among vertebrates includes three months for the east African fish, *Nothobranchius furzeri*, 211 years for the bowhead whale (George, et al. 1999; Valdesalici and Cellerino 2003) and at least 272 years for the Greenland shark (Nielsen, et al. 2016). This range expands further if we include nonvertebrate animals such as the ocean quahog clam (*Arctica islandica*), which can live at least four centuries (Wanamaker Jr, et al. 2008), and the deep-sea oyster *Neopycnodonte zibrowii*, which can live at least five centuries (Wisshak, et al. 2009). As with any species-specific trait, maximum lifespan can be assumed to be at least partially genetic in origin and therefore subject to change through evolution, either via positive selection, negative purifying selection, or through neutral genetic drift. This assumption is supported by the numerous single gene mutations known to extend the lifespan of laboratory mice, fruit flies, and nematodes (Ladiges, et al. 2009), and by the characteristic changes in epigenetic profile and gene expression known to accompany the aging process (De Paoli-Iseppi, et al. 2017; Mayne, et al. 2019; Levine, et al. 2020; Robeck, et al. 2021). Given these aspects of lifespan and aging, it is noteworthy that maximum lifespans can vary dramatically between closely related species (Gorbunova, Bozzella and Seluanov 2008). Furthermore, observations support the possibility that the aging process is intrinsically different between long-lived and short-lived animals (Haussmann, et al. 2003; Copes, et al. 2015). If lifespans and aging are genetically influenced, and closely related species can vary widely in lifespan, then it is reasonable to predict that for some species a relatively small number of genetic changes are required to produce large changes in aging and lifespan, whether these changes are through nonsynonymous mutations or though alterations to gene expression.

The AnAge database is an often-cited source for species maximum lifespans, with currently over 3,700 recorded maximum lifespan estimates (Tacutu, et al. 2013). These estimations are a consensus compiled from a wide variety of sources, including wild and captive species, from large and small samples, under diverse conditions. Accordingly, the quality of the data is categorized by the project curators as *high, acceptable, questionable*, or *low* depending on the source. Comparison between the maximum longevity of any of these species and *Homo sapiens* is somewhat challenging given that our knowledge of human lifespan is based on hundreds of millions of observations, whereas our knowledge of other species is often based on hundreds or thousands of observations at best (Austad 2010). The estimated maximum longevity for any species is partially a function of the number of observations. For example, only about one out of every 10,000 humans lives to the age of 100 (Perls, et al. 2002; Austad 2010). An estimate of human maximum longevity based on the observed lifespan of only 1,000 randomly selected individuals would then likely fall short of 100 years. Therefore, it can be assumed that for species with few observations, the estimated maximum lifespan is likely closer to the average life expectancy than the true maximum for that species. If human life expectancy is used for comparison, AnAge contains *high* or *acceptable* estimates for 45 chordate species with maximum lifespans longer than humans, and 20 with lifespans greater than the human life expectancy upper 95% confidence interval (106.2 years; human life expectancy, plus 1.96 times the standard deviation).

For our study, we examined two chordate species with lifespan estimates greater than this 106.2-year threshold, and with available high-quality gene and protein sequence information – the blue whale (*Balaenoptera musculus*) and the Pinta Island tortoise (*Chelonoidis abingdonii*). Notably, these two animals fall within separate taxonomic classes and are separated evolutionarily by approximately 320 million years. We then identified closely related short-lived animals as points of comparison, using the beluga whale *Delphinapterus leucas* and the Pacific white-sided dolphin *Lagenorhynchus obliquidens* as short-lived relatives of *B. musculus* (both of which live less than 50 years), and the Chinese pond turtle or Reeves’ turtle *Mauremys reevesii* as a short-lived relative of *C. abingdonii* (maximum lifespan: 24.2 years). Investigation of orthologous proteins among these species revealed no evidence of convergent evolution or deleterious effect on phenotype. However, evidence of positive selection was observed in a small set of genes in both long-lived species, as compared to their shorter-lived relatives. The bowhead whale (*Balaena mysticetus*) is a long-lived species, with a maximum longevity of 211 years. This species lacks an annotated genome available through the NCBI, but gene and protein sequence files are available through the Bowhead Whale Genome Resource (http://www.bowhead-whale.org/) (Keane, et al. 2015). After including *B. mysticetus* in our investigation of positive selection, neuregulin-1 (NRG-1) remained as the only ortholog showing evidence of positive selection among all selected long-lived animals.

## Materials and Methods

### Lifespan Categorization

Maximum longevity data was obtained from the publicly available AnAge database (Tacutu, et al. 2013), which contains a curated list of maximum longevity estimates for 4,244 species (build 14). The AnAge dataset is constructed both from animals observed in the wild and from those maintained in captivity, and the estimates are ranked based on the quality of the supporting evidence (ranked either as *high, acceptable, low*, or *questionable*). We further categorized maximum longevity estimates based on comparison to the worldwide average human life expectancy. Using an average human life expectancy past age ten of 77.4 years, ± 14.7 years standard deviation (Edwards 2013), we calculated the 95% confidence interval for human life expectancy as the range from 48.6 years to 106.2 years (+/- 1.96 × 14.7 years). Animals with a reported maximum longevity beyond this 106.2-year cutoff were categorized as *long-lived*. Using this same approach, *short-lived* animals were defined as those with a reported maximum longevity below the 48.6-year cutoff, and *midrange* lifespan was defined as falling within the inclusive 48.6-year to 106.2-year range. This categorization scheme is admittedly less than ideal given that humans fall into the long-lived category when including the many known supercentenarians, and especially when including the recorded human maximum longevity of 122.5 years (Robin-Champigneul 2020). This categorization scheme also has the additional weakness of being based on the comparison of maximum animal longevity to average human life expectancy, which are two different types of measures. Additionally, comparison between the maximum longevity of any of these species and *Homo sapiens* is somewhat challenging given that our knowledge of human lifespan is based on hundreds of millions of observations, whereas our knowledge of other species is often based on hundreds or thousands of observations at best (Austad 2010). The estimated maximum longevity for any species is partially a function of the number of observations. For example, only about one out of every 10,000 humans lives to the age of 100 (Perls, et al. 2002; Austad 2010). An estimate of human maximum longevity based on the observed lifespan of only 1,000 randomly selected individuals would then likely fall short of 100 years. Therefore, it can be assumed that for species with few observations, the estimated maximum lifespan is likely closer to the average life expectancy than the true maximum for that species. In other words, examples of extreme human longevity are assumed to be over-represented in the available observations while species-specific examples of extreme animal longevity are likely underrepresented, which implies that average human life expectancy and reported animal maximum longevity should be closely comparable in many instances.

### Selection of Species for Analysis

Animals were selected for initial analysis based on the criteria of having a maximum longevity estimate available from AnAge ranked as *high* or *acceptable*, and having an annotated reference genome available through the National Center for Biotechnology Information (NCBI). To be included, an animal also had to be categorized as follows: (1) long-lived, or (2) short-lived and the closest available phylogenetic relative to a long-lived animal. To find available candidates, the AnAge database was cross-referenced with evolutionary divergence times found in the TimeTree database (Hedges, Dudley and Kumar 2006; Kumar and Hedges 2011; Hedges, et al. 2015; Kumar, Stecher, et al. 2017). The NCBI Entrez Programming Utilities API was accessed to find FTP links for downloading eukaryotic reference genome files. All files were downloaded January 7^th^, 2021. Among the long-lived species, only seven currently have genome files available from the NCBI and only two have protein sequence files (*Balaenoptera musculus* and *Balaenoptera physalus*). Of these two species, only the blue whale, *Balaenoptera musculus*, has an annotated reference genome available from NCBI. To expand the list of species for possible analysis, a literature search was performed, which revealed the Pinta Island Tortoise, *Chelonoidis abingdonii*, as has having a reference sequence and protein sequence files available from the NCBI, and a maximum longevity estimated to overlap the 106.2-year threshold (100-years to 120-years) (Mayne, et al. 2019; Quesada, et al. 2019). Short-lived relatives for *B. musculus* and *C. abingdonii* were selected following a similar methodology. The TimeTree database was used to find species that (1) share a most recent common ancestor (MRCA) with *B. musculus* and *C. abingdonii* within the last 100 million years (Hedges, Dudley and Kumar 2006; Kumar and Hedges 2011; Hedges, et al. 2015; Kumar, Stecher, et al. 2017), (2) have a *high* or *acceptable* quality maximum longevity less than 48.6 years (less than the lower range of the 95% CI for human life expectancy) as reported in the AnAge database, and (3) have a reference sequence and protein file available from NCBI. Of these species, the closest relatives were chosen based on the identified time to the most recent common ancestor (TMRCA; Table 1) and consist of *Delphinapterus leucas* (beluga whale) and *Lagenorhynchus obliquidens* (Pacific white-sided dolphin) as the most-closely related short-lived species to *B. musculus*, and *Mauremys reevesii* (Chinese pond turtle or Reeves’ turtle) as the most-closely related short-lived species to *C. abingdonii*. To further investigate orthologs with evidence of positive selection, one additional long-lived species was identified for analysis. The bowhead whale (*Balaena mysticetus*) has a reported lifespan of 211 years (George, et al. 1999), with a CpG dinucleotide estimated lifespan of above the 205-year calibration range of the model (267.9 years; see “Lifespan Estimation Using Gene Promoter CpG Density” below). *B. mysticetus* currently lacks inclusion among the NCBI databases, however protein and gene sequence files were downloaded from The Bowhead Whale Genome Resource (http://www.bowhead-whale.org/) (Keane, et al. 2015).

**Table 1:**
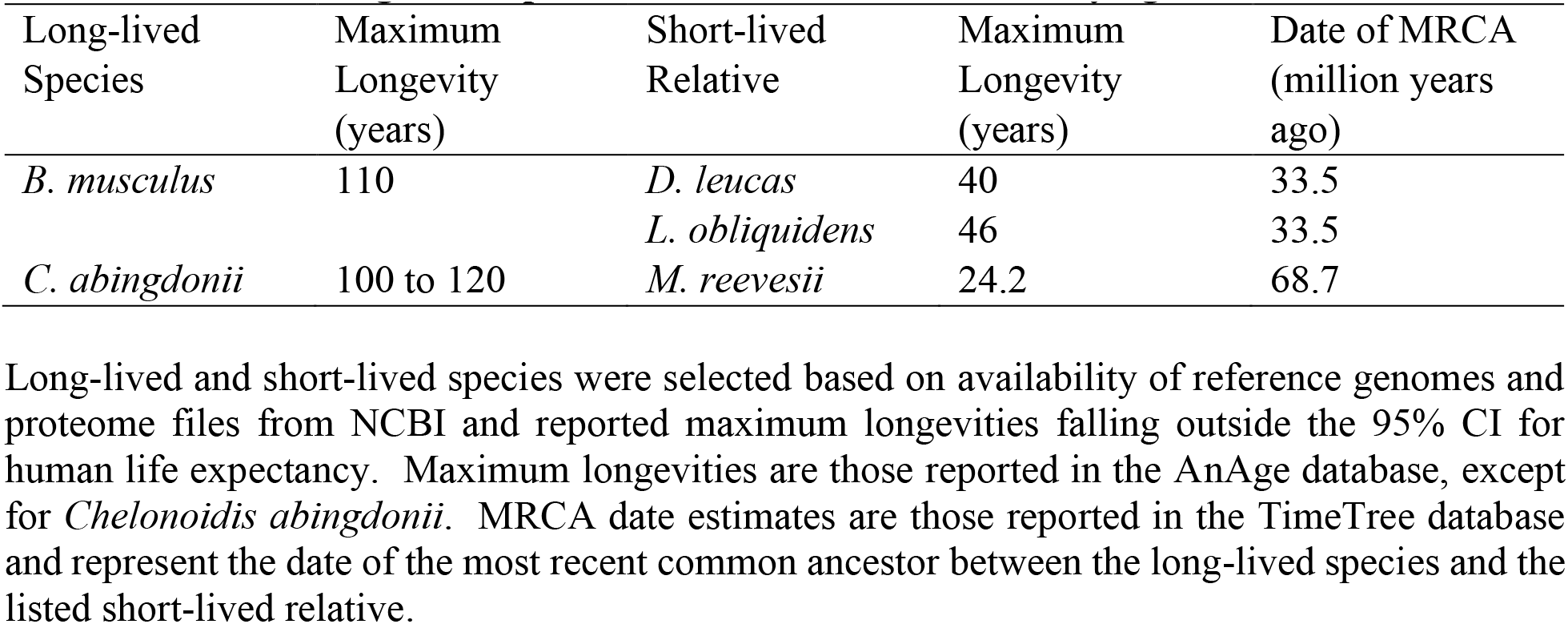
Selected Long-lived Species and Closest Short-lived Phylogenetic Relatives.

| Long-lived<br>Species | Maximum<br>Longevity<br>(years) | Short-lived<br>Relative | Maximum<br>Longevity<br>(years) | Date of MRCA<br>(million years<br>ago) |
| --- | --- | --- | --- | --- |
| *B. musculus* | 110 | *D. leucas* | 40 | 33.5 |
|  |  | *L. obliquidens* | 46 | 33.5 |
| *C. abingdonii* | 100 to 120 | *M. reevesii* | 24.2 | 68.7 |

Long-lived and short-lived species were selected based on availability of reference genomes and proteome files from NCBI and reported maximum longevity falling outside the 95% CI for human life expectancy. Maximum longevity are those reported in the AnAge database, except for *Chelonoidis abingdonii*. MRCA date estimates are those reported in the TimeTree database and represent the date of the most recent common ancestor between the long-lived species and the listed short-lived relative.

### Lifespan Estimation Using Gene Promoter CpG Density

Lifespan estimation based on genetic sequence was performed using the method published by Dr. Benjamin Mayne (Mayne, et al. 2019). In summary, the CpG density within 42 individual gene promotor sequences was reported to correspond to vertebrate lifespan within taxonomic classes. The region -499 to +100 nt for each promoter was queried against the selected genome using BLAST v2.10.1 (Altschul, et al. 1997) The following command parameters were used:

blastn -query <human promoters.fasta> -db <target genome> -task megablast -max_hsps 1 - outfmt “6 qseqid qlen qstart qend sacc sstart send evalue bitscore length pident qcovhsp qseq sseq” -culling_limit 1 > <output file>

For each high-scoring segment pair (HSP), CpG density was calculated as the number of CpG dinucleotides identified within the corresponding region of the subject sequence, divided by the length of the HSP. The *raw density* was then calculated as follows:

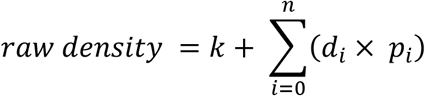

Where *k* is an intercept value, *n* is the number of HSPs identified, *d* is the CpG density, and *p* is a unique coefficient associated with each gene promoter. Using the raw density, lifespan was estimated as:

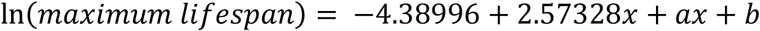

Where *x* is the raw density, and *a* and *b* are coefficients associated with taxonomic class (for *Reptilia, a* = -0.48958 and *b* = 1.17281; for *Mammalia, a* = -0.92888 and *b* = 2.33508). For long-lived non-mammalian vertebrates, the maximum lifespan value was then multiplied by *x*.

### Lifespan Estimation Using Allometry

Adult body mass has been reported to positively correlate with vertebrate lifespan across taxonomic groups (Promislow 1993; de Magalhães, Costa and Church 2007). To calculate estimated maximum lifespan based on body mass, the allometric equation reported by de Magalhães et. al. (2007) for mammals, birds, reptiles, and amphibians was used as follows:

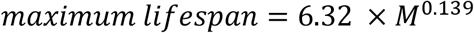

Where *M* is the species adult body mass in grams (*r*^2^ = 0.40 for correlation with lifespan). This equation was used to estimate maximum lifespan for *Balaenoptera musculus, Delphinapterus leucas*, and *Lagenorhynchus obliquidens* given that the other equations for mammals reported by the authors specifically exclude cetaceans. For *Chelonoidis abingdonii* and *Mauremys reevesii* the following reptile-specific equation was used (*r*^2^ = 0.35):

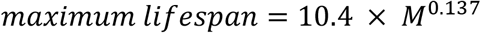

The longevity quotient (LQ) was also calculated for all species as the reported lifespan divided by the allometrically estimated lifespan (Yu, et al. 2021). This quotient provides a measure of longevity scaled to body mass, with values above or below 1.0 being long-lived or short-lived respectively, relative to the body mass. Adult body mass was obtained from AnAge for *B. musculus, D. leucas*, and *L. obliquidens*, and from the reptile body size dataset from Gorman and Hone et. al. 2012 for *C. abingdonii* and *M. reevesii* (O’Gorman and Hone 2012).

### Identification of Orthologs

For identifying protein sequence orthologs, top scoring reciprocal best hits (RBHs) were found between all pairs of species using protein sequence files and BLAST v2.10.1 (Altschul, et al. 1997; Tatusov, Koonin and Lipman 1997; Bork, et al. 1998; Ward and Moreno-Hagelsieb 2014) using the following parameters:

blastp -query <query species.fasta> -db <target proteome.fasta> -outfmt “6 qaccver saccver salltitles” -out <output file> -matrix <substitution matrix> -evalue 1e-6 -max_hsps 1 - num_alignments 1 -num_threads 19 -soft_masking true -subject_besthit -use_sw_tback For the BLAST searches, the substitution matrix was chosen based on the evolutionary divergence times between the species pairs as reported in the TimeTree database (Hedges, Dudley and Kumar 2006; Kumar and Hedges 2011; Hedges, et al. 2015; Kumar, Stecher, et al. 2017), and as previously suggested for maintaining an acceptable balance between BLAST sensitivity and HSP length (Pearson 2013). BLOSUM80 was used for pairs of species that diverged by 200 million years or less, and BLOSUM62 was used for species that diverged by more than 200 million years. For each pair of species, the BLAST search was performed twice, switching the query and subject species FASTA files for the second search. RBHs were then identified by using a custom JavaScript script to search both BLAST output files for proper matches. Proteins were determined to be orthologs if: (1) their accession numbers were exactly matched to HSPs within both output files, or (2) they were matched to isoforms of the same protein, as determined by NCBI FASTA file deflines.

### Graph Analysis of Homologs

Semi-global protein sequence alignments were generated for each RBH using BLOSUM62 with a gap open penalty of 11 and a gap extension penalty of 1 for all internal gaps. Where homologs involved isoforms of the same protein, the longest sequence isoform from both species was used for the alignment. The score, query sequence, subject sequence, accession numbers, and deflines for each alignment were then saved as output for each pair of species. Sets of orthologous proteins were selected for further analysis by constructing graphs representing shared homologies. For this process, separate proteins were represented as graph nodes. Each semi-global alignment between homologous proteins pairs was then represented as a line connecting those two nodes. A set of nodes was determined to be valid if (1) a continuous path could be traced through one node from each species by following the connecting lines, and (2) each node was directly linked to all other nodes within that set by one line. Using this method allowed for individual nodes to participate in more than one valid set, and for each set to consist exclusively of mutually homologous proteins.

To investigate possible protein sequence-level similarities between the two long-lived species, orthologous sets were selected based on having higher protein sequence alignment scores between the two long-lived species (*B. musculus* and *C. abingdonii*) than between them and any of the short-lived species (i.e., higher scores along line *i* of the graph in Supplemental Figure S1 panel A than along lines *ii, iii*, and *v*). Similarity ratios were also calculated for each orthologous set dividing the alignment score for the two long-lived species (score along line *i*) by the alignment score for a long-lived species and a closely related short-lived species (score along lines *ii, iii*, and *v*). Using this method, three ratios were calculated for each set of orthologs. When calculated for all orthologous sets, ratios exhibited a Pareto distribution as determined by the fitdistrplus package in R. Estimation of distribution shape and scale permitted the calculation of associated *p-values*. Multiple comparisons were corrected using the Benjamini-Hochberg method (*p* < 0.05; FDR < 0.1), and orthologs with ratios greater than the threshold were excluded. A similar methodology was used to investigate the selected short-lived animals (i.e., identification of scores higher along lines *v, ix*, and *x* than along lines *ii, iii*, and *v*). For the short-lived species, any orthologs that were not identified in all comparisons were excluded (see Supplemental Table S1 for a complete high-scoring ortholog list).

### Tree Production and Visualization

For initial analysis of the entire array of graphs (sets of orthologs), a median score was calculated between each pair of species from the homologous protein sets. Unweighted pair group method with arithmetic mean (UPGMA) was then used to construct a phylogenetic tree, with the negative median scores used as a substitution for distance. Visualization of trees was performed by uploading Newick files to Tree Viewer (http://etetoolkit.org/treeview/) (Heurta-Cepas, Serra and Bork 2016).

### Identification of Positively Selected Genes

For selected sets of orthologs, protein multiple sequence alignments were generated using T-Coffee, version 13.45.0.4846264 (Notredame, Higgins and Heringa 2000), and nucleotide alignments were generated from the resulting files using the nucleotide sequences downloaded from the NCBI and PAL2NAL version 14 (Suyama, Torrents and Bork 2006). Positive selection was tested using PAML version 4.9 (Yang 2007) and the phylogenetic tree for the five selected species generated from the TimeTree database (Supplemental Figure S1 panel C). The branch site model for codon evolution was used (model = 2; NSites = 2). For each analyzed set of orthologs, a model allowing for positive selection of codons (fix_omega = 0) was compared to the null model of neutral or purifying selection (fix_omega = 1; omega = 1). Likelihood ratio tests (LRTs) were performed by PAML, and comparison of LRTs was calculated using a χ^2^ test with an assumed 50:50 distribution and a point mass at zero. Significance was determined after correcting for multiple comparisons using the Benjamini-Hochberg method (*p* < 0.05) with a false discovery rate (FDR) cutoff of 0.1. Positive selection was tested seven different times for each selected set of orthologs; once with the long-lived species selected as the foreground branches; once with the short-lived species selected as the foreground branches; and once with each of the five separate species selected as a foreground branch. Orthologs were considered for further analysis if they displayed positive selection with the long-lived species (separately and as a group), or with the short-lived species (separately and as a group). Given that phylogenetic tree discordance can affect the ability of PAML to detect positive selection (Mendes and Hahn 2016), MrBayes version 3.2.6 (Ronquist and Huelsenbeck 2003) was used to generate a phylogenetic tree for each selected set of orthologs (lset nst=6 rates = invgamma; mcmc ngen=10000 samplefreq=10; sump burnin=250). The PAML positive selection tests were then performed again using the tree obtained from MrBayes when the topology of the phylogenetic tree differed from the one obtained from TimeTree. Orthologs were excluded from further analysis if they did not display positive selection using the phylogenetic tree generated from MrBayes.

For eukaryotic organisms, species-specific isoforms and the incomplete identification of isoforms can both lead to alignment of nonhomologous regions. These regions of misalignment then risk being interpreted as evidence of evolution during dN/dS analysis, increasing the likelihood of false positives (Villanueva-Cañas, Laurie and Albà 2013). To compensate for this risk, orthologs with evidence of positive selection under all the above conditions were subjected to manual inspection. Isoforms were examined from each species, and if an alternative presented a better alignment it was selected, otherwise the longest isoform was retained. New multiple sequence alignments were generated where appropriate, and the alignments were inspected and manually adjusted where it was deemed necessary. Orthologs that were updated by manual inspection were then re-subjected to the testing procedure outlined above.

### Prediction of Deleterious Proteins

Multiple sequence alignments generated from T-Coffee (see above) were analyzed to identify sequence locations where either (1) the long-lived species sequences were identical and different from any short-lived species at that sequence location, or (2) the short-lived species sequences were identical and different from any long-lived species at that sequence location. For these qualifying locations, files were generated that contained comma separated variant information for each species. Protein FASTA files and variant files were then supplied to PROVEAN version 1.1 for estimation of phylogeny-corrected effect prediction (Choi, et al. 2012). PROVEAN is designed for binary classification of sequence variants as either *neutral* or *deleterious*, with scores below -2.282 providing a 79% accuracy in the classification of deleterious proteins against a UniProt human variant dataset. For the selected ortholog sets, a conservative threshold score of -5.0 was chosen for classification given its previous successful use in species-to-species comparisons (Burga, et al. 2017). Variants were considered deleterious if they produced a score below this threshold in all species examined.

### Gene Ontology

Over-representation analysis of gene ontology terms and KEGG pathways was performed using GeneTrail2 (Stöckel, et al. 2016), with upper-tailed hypothesis testing and Benjamini-Hochberg multiple comparison correction (*p* < 0.05).

## Results

### Independent estimation of lifespan for selected species

CpG dinucleotide estimation of maximum lifespan for the chosen species corresponded closely to those reported by AnAge (*r*^2^ = 0.818; *p-value* = 0.063; Table 2) except for *B. musculus*, which produced an estimate greater than the 205-year upper calibration range used in the original method. The CpG dinucleotide estimation method also over-estimated the lifespan for *M. reevesii* as 52.5 years, placing this species in the *midrange* lifespan category defined above (48.6 years to 106.2 years). Allometric lifespan estimation based on body mass underestimated maximum longevity for *B. musculus, L. obliquidens*, and *C. abingdonii*, and this underestimation is reflected in the longevity quotient (LQ) measures for these three species. The LQ results for these three species suggest a greater lifespan than expected from body mass alone. However, an analysis of 987 mammalian species by Yu et. al. (2021) provides a standard deviation of ± 0.57 for mammal LQ, which places the LQ for *B. musculus* and *L. obliquidens* within the usual variation observed for this taxonomic group.

**Table 2:**
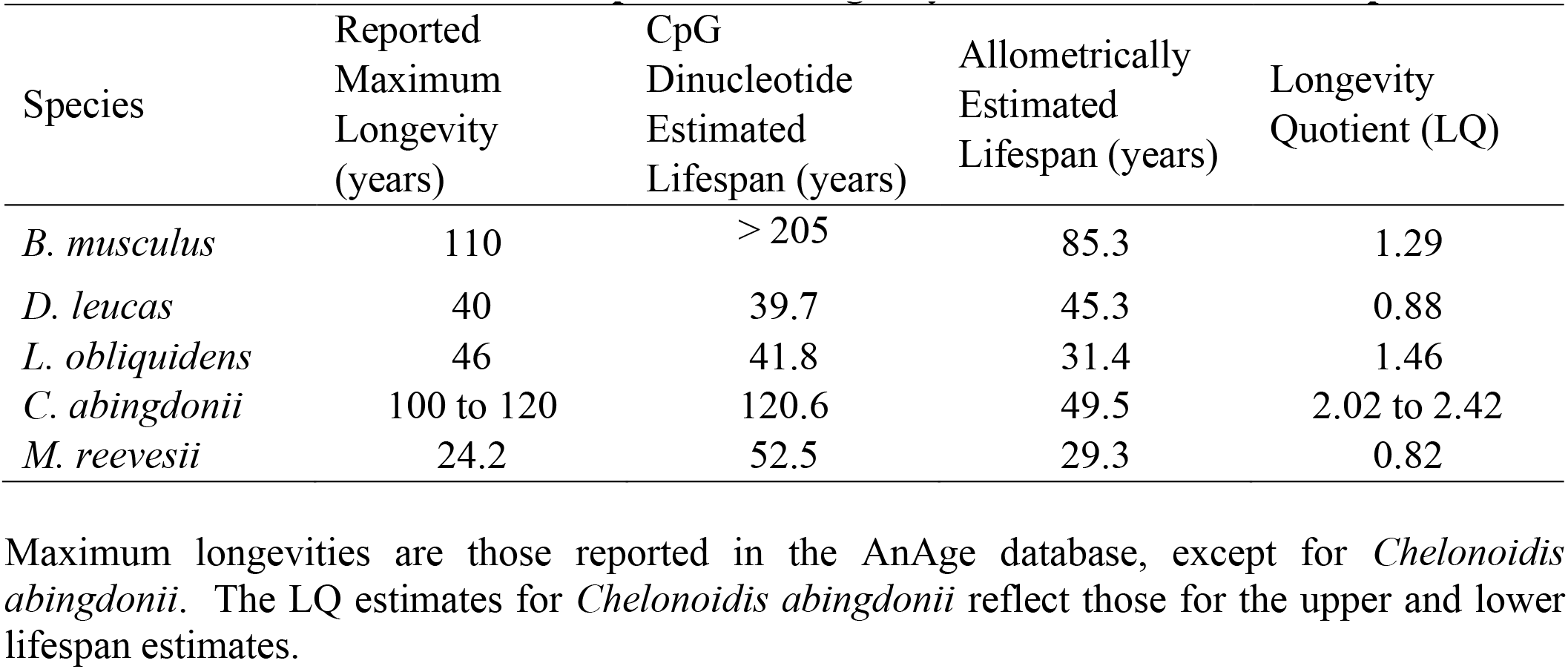
Estimated Maximum Lifespans and Longevity Quotients for Selected Species.

### Orthologous protein alignment

The pattern of alignment scores matched the known evolutionary relationship among these species, with higher median scores present between the two reptiles (*C. abingdonii* and *M. reevesii*) and among the three Cetaceans (*B. musculus, D. leucas*, and *L. obliquidens*). Median alignment scores also closely correlated with TMRCA values found in the TimeTree database (*r*^2^ = 0.997; p-value = 2.485 × 10^−11^). Since each set of orthologous proteins contains an all-to-all pairwise sequence alignment, we were able to generate a phylogenetic tree for these species using UPGMA and directly utilizing median scores as a distance metric (Supplemental Figure S1 panel B). The resulting tree matches closely with the phylogenetic tree for these species produced by the TimeTree database (Supplemental Figure S1 panel C) and demonstrates that the distribution of protein alignment scores among the selected species reflects their overall evolutionary relationship. In total, 10 orthologs were identified as having a higher alignment score between long-lived species (*B. musculus* and *C. abingdonii*), and 21 orthologs were identified as having a higher alignment score between short-lived species (*D. leucas, M. reevesii*, and *L. obliquidens*). Two orthologs appeared in both sets, actin-related protein 2 (ACTR2) and protein transport protein Sec61 subunit alpha (SEC61A), giving a total of 29 unique proteins among the two sets (Supplemental Table S1).

Over-represented gene ontology terms for these high-scoring proteins include ErbB signaling (*p* = 0.0232) and ubiquitin mediated proteolysis (*p* = 0.0232) among the long-lived species, and acetylgalactosaminyltransferase activity (*p* = 0.0418) and ribosome binding (*p* = 0.0418) among the short-lived species. Investigation of the complete list of 29 orthologs was enriched in terms including PI3K/AKT activation (*p* = 0.0434), cellular response to hypoxia (*p* = 0.0031), cellular responses to stress (*p* = 0.0182), adaptive immune system (*p* = 9.02 × 10^−6^), Toll-like receptors cascades (*p* = 0.0453), class I MHC mediated antigen processing (*p* = 0.0016), and DNA methylation (*p* = 0.0464; see Supplemental Table S2).

### Deleterious effect on phenotype

Analysis of the multiple sequence alignments for each of these orthologs did reveal a series of protein sequence locations in three of the 29 orthologs that were either (1) identical at that location in both long-lived species and different at that location from the short-lived species, or (2) identical in all short-lived species and different at that location from either of the long-lived species sequences. Identification of these locations generated a series of sequence features that were consistent within each lifespan category. PROVEAN version 1.1 was used to predict possible functional effects of these sequence features (Choi, et al. 2012). For each of the three orthologs with characteristic long-lived or short-lived sequence features, we examined the predicted impact of applying that feature as a sequence variant across the selected species. Using the conservative score threshold of -5.0, all identified sequence features were reported as neutral.

### Positive selection and convergent evolution

Among these 29 orthologs, evidence of possible positive evolutionary selection was investigated using dN/dS analysis, and the software PAML version 4.9 (Yang 2007). Three orthologs were found with statistically significant evidence of positive selection reported by PAML in tested foreground branches, consisting of NRG-1, GALNT17, and SCFD1. Pro-neuregulin-1, membrane-bound isoform (NRG-1) and N-acetylgalactosaminyltransferase 17 (GALNT17) were found to have significant positive selection exclusively among the long-lived species, while Sec1 family domain-containing protein 1 (SCFD1) was positively selected among all species examined except for *L. obliquidens* (Supplemental Table S3). One additional ortholog, phospholipid phosphatase-related protein type 1 (PLPPR1), was found to have partial evidence, but produced a *p*-value short of the multiple comparison corrected threshold when using *B. musculus* as a foreground branch in the analysis (*p* = 0.181). The possibility of sequence-level convergent evolution was investigated, however none of the sequence locations reported by the PAML naïve empirical Bayes algorithm with a posterior probability above 95% were identical in either the long-lived or short-lived species exclusively.

Fewer protein sequences are available for *B. mysticetus* compared to the other species used (e.g., 22,672 sequences for *B. mysticetus* versus 52,259 for *B. musculus*). Of the three orthologs with evidence of positive selection, GALNT17 was absent among the *B. mysticetus* sequences, leaving only NRG-1 and SCFD1 for analysis by PAML. Using *B. mysticetus* as a foreground branch in the analysis generated evidence of positive selection only for NRG-1.

## Discussion

Using PAML, we were able to detect evidence of positive selection in three of the examined orthologs, two of which (NRG-1 and GALNT17) showed evidence of positive selection exclusively with the long-lived species *B. musculus* and *C. abingdonii*. Notably, NRG-1 and GALNT17 are both associated with neural development and function (Mei and Xiong 2008; Weisner, et al. 2019). NRG-1 acts in vivo as a direct ligand for ERBB3 and ERBB4, members of the epidermal growth factor receptor (EGFR) family of receptor tyrosine kinases. Variants of NRG-1 have been identified as possible risk factors for schizophrenia (Munafo, et al. 2006; Mei and Xiong 2008), and altered expression of NRG-1 in the brain appears to play a role in the pathophysiology of Alzheimer’s disease (Jiang, et al. 2016; Mouton-Liger, et al. 2020). GALNT17 is less well characterized, but it is expressed in multiple cell types in the brain and serves as an N-acetylgalactosaminyltransferase that ultimately affects cell adhesion and motility, pinocytosis, and lamellipodia formation (Nakayama, et al. 2012). Deletion of GALNT17 occurs in Williams-Beuren syndrome, and likely contributes to the associated phenotype (Merla, et al. 2002; Weisner, et al. 2019).

Of the two proteins with complete and significant evidence of positive selection among the long-lived animals, only NRG-1 has an identifiable ortholog in the current assembly of the bowhead whale genome. In total, NRG-1 displayed evidence of positive selection for all three long-lived animals (*B. musculus, B. mysticetus*, and *C. abingdonii*). No evidence was found suggesting convergent evolution for NRG-1 among these species, and no shared deleterious sequence features were predicted between *B. musculus* and *C. abingdonii*. No claim can be made here regarding the underlying source of positive selection, and this identification of positive selection in no way directly connects NRG-1 to the longevity of these species. However, it is noteworthy that intracellular NRG-1 signaling has been demonstrated to be neuroprotective following brain ischemia (Navarro-González, Huerga-Gómez and Fazzari 2019), and NGR-1 was identified as a candidate human longevity gene in a study examining variants found among exceptionally long-lived siblings (Cash, et al. 2014). The three long-lived animals examined in our study are a diverse group, including a terrestrial reptile and two aquatic mammals from separate genera. Besides an extended lifespan, all three species are physically larger than the short-lived animals we used for comparison, which highlights the possibility that positive selection of NRG-1, GALNT17, and PLPPR1 could be a signature of enlarged brain volume. It is especially suggestive then that cerebellar expression of NRG-1 has been found to positively correlate with maximum lifespan among multiple rodent species independent of brain mass or body size (Edrey, et al. 2012).

