## Supplementary material for "Positive selection of neuregulin 1 (NRG-1) among three long-lived vertebrates": Figure S1

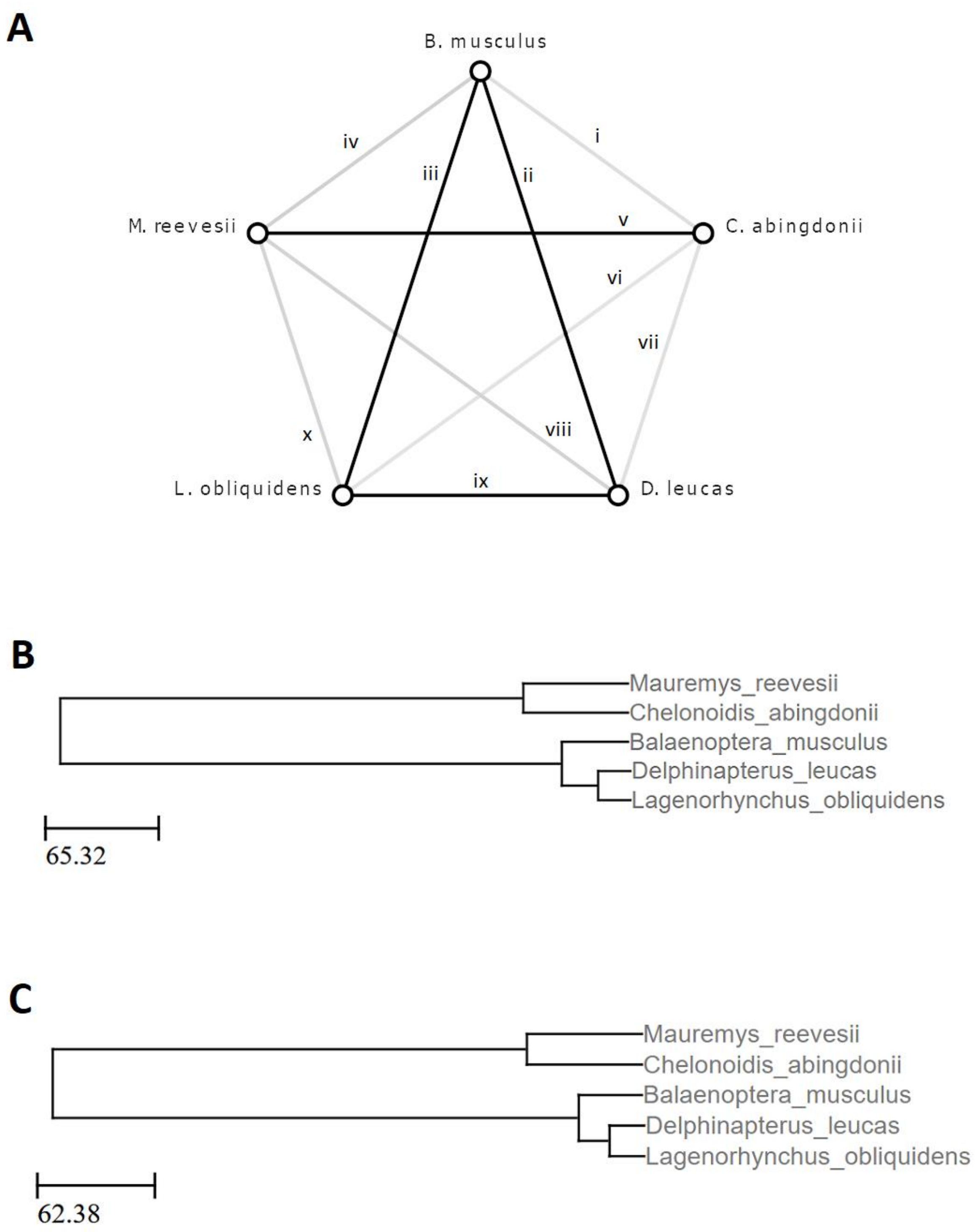


**Figure 1: Reconstructed Evolutionary Relationship Among Selected Species.** (A) Lines represent the median semi-global alignment scores. Darker lines indicate a higher median score. Lines are numbered for reference. (B) Tree produced from UPGMA using negative median score in substitution of a distance metric. The Lines have been scaled using the equation of linear

regression found between the negative median score and the TMRCA found in the TimeTree database ($y = 0.3913x + 995.41$; where $y$ is the time in millions of years, and $x$ is the median score). (C) The phylogenetic tree downloaded from the TimeTree database.
